# CRISPR-Cas9 HDR Optimization: RAD52, Denatured and 5’-Modified DNA Templates in Knock-In Mice Generation

**DOI:** 10.1101/2025.05.05.652162

**Authors:** Boris V. Skryabin, Daniela A. Braun, Helena Kaiser, Leonid Gubar, Birte Seeger, Tasneem Khanam, Anja Stegemann, Hermann Pavenstädt, Timofey S. Rozhdestvensky

## Abstract

CRISPR/Cas9-mediated genome editing is a powerful tool for producing animal models of human diseases. However, it often encounters challenges related to low efficiency of donor DNA templates insertion through homology-directed repair (HDR) pathway or unwanted insertions and/or multiplications. Here, we present findings from multiple targeting experiments aimed at generating a Nup93 conditional knockout (cKO) mouse model. Injection of CRISPR/Cas9 components into over two thousand zygotes, resulted in 270 founder animals. Our study revealed various obstacles associated with the use of single-stranded (ssDNA) and double-stranded DNA (dsDNA) templates during cKO generation, highlighting the critical role of denaturation of long 5’-monophosphorylated dsDNA templates in enhancing precise genome editing and reducing template multiplications. Application of RAD52 protein increased HDR efficiency of ssDNA integration almost 4-fold, albeit with an associated increase in template multiplication. Targeting the antisense strand of DNA using two crRNAs demonstrated better efficacy in HDR-mediated precise genome editing when compared to targeting the sense or sense-antisense strands. In addition, the application of 5’-end biotin-modified donor DNA resulted in up to a 8-fold increase in HDR-mediated single-copy template integration compered to unmodified dsDNA donor. Furthermore, application of 5’-end C3 spacer modified template resulted in up to a 20-fold increase in correctly HDR modified mice independent from ssDNA or dsDNA template employment. This study underscores potential pitfalls in CRISPR/Cas9-mediated genome editing and offers simple practical solutions to refine this potent tool. These findings highlight various strategies to enhance CRISPR/Cas9 HDR efficiency, providing a framework for improving precision in the generation of conditional knockout models.

## Introduction

Generating gene knock-in (KI) and conditional gene knockout (cKO) animal models are among the most complex and time-consuming objectives in the production of genetically engineered species. Site-specific recombinase (SSR) systems such as Cre-LoxP from bacteriophage P1 and Flp-FRT from Saccharomyces cerevisiae are commonly utilized in creating cKO mouse models. In generating cKO mice, critical exons of the target gene are flanked by directly oriented LoxP or FRT sites. The Cre or Flp recombinase recognizes and catalyses the deletion of the DNA region between these sites, respectively [1]. Despite certain technical limitations, cKO is considered one of the most reliable and sophisticated approaches for investigating gene function *in vivo*. It allows researchers to overcome issues of embryonic or early postnatal lethality observed in conventional knockout animals and enables the study of gene function in a tissue- or cell-type-specific manner [1–4].

Nowadays, CRISPR/Cas9 endonuclease-mediated genome editing has revolutionized biological and biomedical sciences. The CRISPR/Cas9 system is a ribonucleoprotein complex comprising two RNA molecules: crRNA (CRISPR RNA) and tracrRNA (trans-activator RNA), which bind to the Cas9 endonuclease [5–7]. The crRNA guides the Cas9 to a specific 20-nucleotide DNA sequence, while the tracrRNA facilitates the assembly of the complex. Cas9 cleaves the double-stranded DNA (dsDNA) approximately 3 bp upstream of the PAM motif (NGG-sequence) [8, 9]. The resulting dsDNA breaks are repaired by either non-homologous end joining (NHEJ) or HDR mechanisms [10]. NHEJ is typically used to create conventional knockout models across a wide range of organisms, whereas HDR enables precise genome editing, including point mutations, in-frame epitope additions, and specific gene knock-ins [11–14]. Efficient integration of large (up to 2000 bp) dsDNA donor fragments into the mouse genome via HDR using CRISPR/Cas9 endonuclease has been demonstrated, however, this process is rare due to the dominance of error-prone nonhomologous end joining (NHEJ), leading to insertions or deletions. Alternative strategies, such as introducing 3′-overhang dsDNA donors or ssDNA in circular or linear formats, CRIS-PITCh, Easi-CRISPR, and others, leverage different donor designs and repair mechanisms [15–21]. For instance, CRIS-PITCh employs short homology arms via microhomology-mediated end joining MMEJ, while Easi-CRISPR uses long single-stranded donors, achieving knock-in efficiencies of up to 60% or higher. We have previously reported that linear dsDNA often multimerizes and integrates as concatemers, complicating precision editing [22].

Recent studies show that 5′ modifications, like biotinylation, can reduce multimerization and improve single-copy HDR efficiency [23–26]. For example, using biotinylated donors with Cas9- Streptavidin fusion proteins increased HDR efficiency up to four-fold, attributed to enhanced recruitment of the donor to the Cas9 complex [27].

Here we investigated different parameters affecting the efficiency of mouse genome editing by CRISPR/Cas9. We perform a comprehensive analysis of different approaches in *Nup93* locus targeting using a double-stranded DNA (dsDNA) and denatured DNA (single-stranded DNA (ssDNA)) templates, showing the critical role of denaturation of long 5’-monophosphorylated dsDNA templates in enhancing precise genome editing and reducing template multiplications. We also have investigated the results of application of RAD52 protein on HDR efficiency of ssDNA integration. In addition, we investigated application of 5’-end C3 spacer (5’-SpC3 or 5’-Propyl) and biotin-modified ssDNA and dsDNA donor to enhance HDR-mediated single-copy template integration.

## Results

### Nup93 locus targeting strategy

To target the *Nup93* genomic locus, we employed a one-step strategy to generate conditional knockout (cKO) mouse models using CRISPR/Cas9 complexes and long donor DNA templates containing two *LoxP* sites. Computational analysis predicted that deletion of exon 9 would introduce a translational frameshift, leading to *Nup93* gene inactivation. Accordingly, we designed a donor DNA fragment with *LoxP* sites flanking exon 9 of the *Nup93* gene (Figure 1A-C, Figure 2). Our one-step insertion strategy relied on the cellular homology-directed repair (HDR) mechanism.

**Figure 1.**
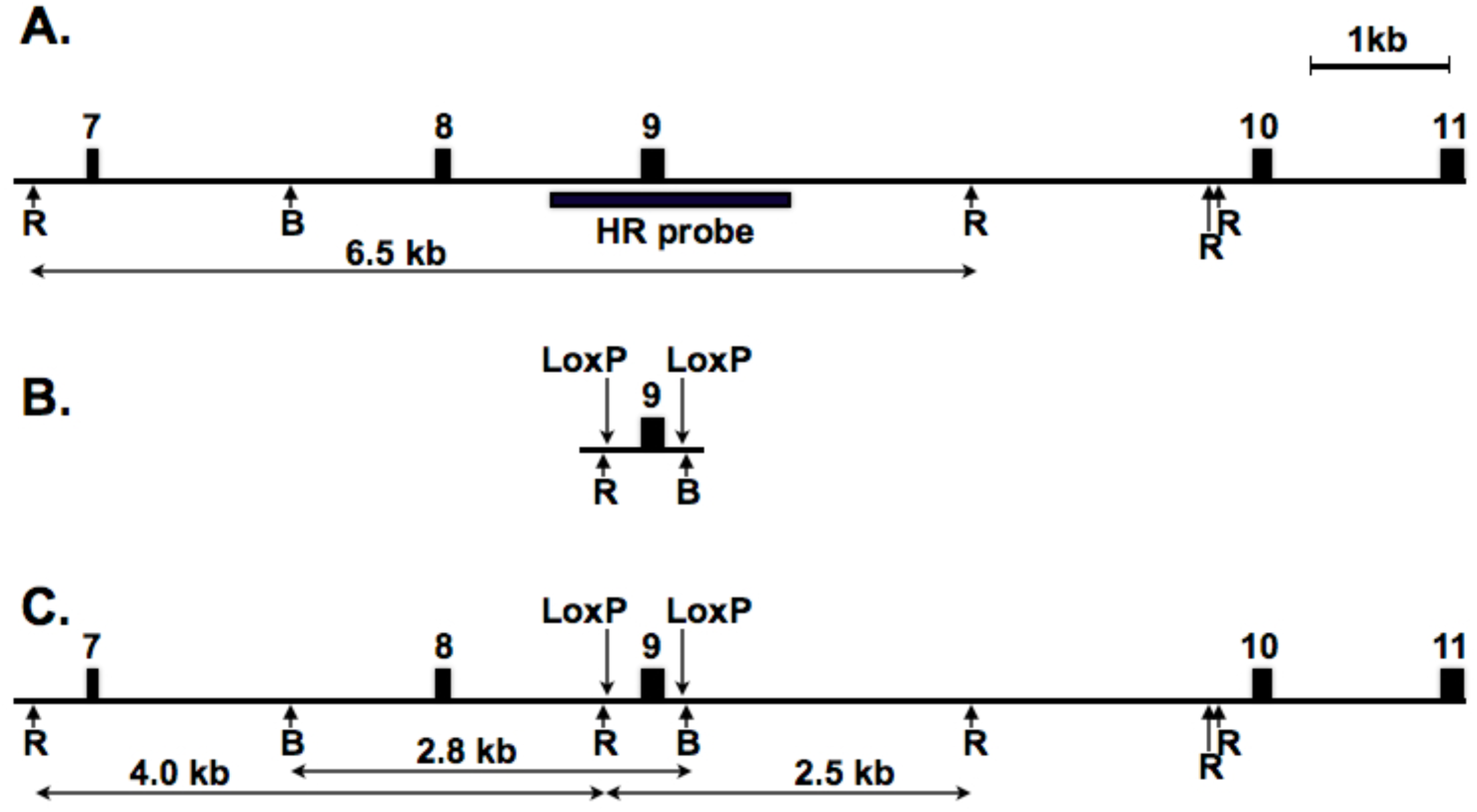
Targeting strategy of the exon 9 of mouse *Nup93* gene**. A-C**: The intron regions are shown as line, exons are shown as filled boxes numbered above. The vertical arrows denote the two *Lox*P sites, and arrows below indicate restriction endonuclease sites *Eco*RI (R) and *Bam*HI (B). The horizontal black bars designated as “HR probe” correspond to area hybridized with donor DNA specific probe used for Southern probe analysis. The expected sizes of restriction DNA fragments are labeled below in kb. **A**: Wild type locus. **B**: Schematic representation of *Nup93* donor DNA template. **C**: Genomic locus after the homologous recombination.

**Figure 2.**
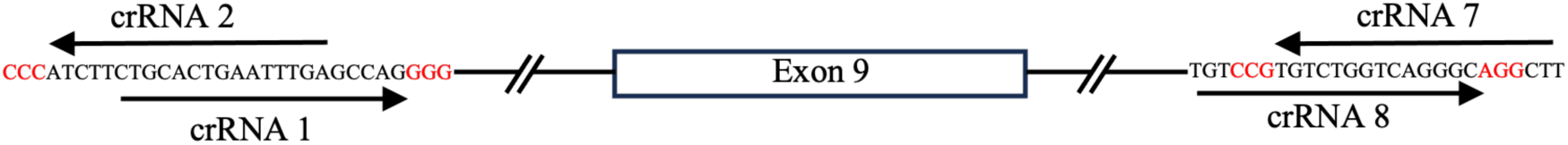
Location of selected crRNAs in the intronic regions flanking exon 9 of the mouse *Nup93* gene. Intronic regions are represented by lines, while exon 9 is depicted as a rectangle. The exact sequences of the crRNA target sites are shown, with PAM sites highlighted in red. Horizontal black arrows indicate the crRNA sequences and their respective orientations (forward “+” or reverse “-”) relative to the annotated *Nup93* gene sequence (Table 1). The target strand refers to the DNA strand complementary to the crRNA, which is cleaved by the HNH domain of the Cas9 protein. Therefore, a crRNA with a “+” orientation targets the antisense DNA strand, and vice versa.

**Table 1.**
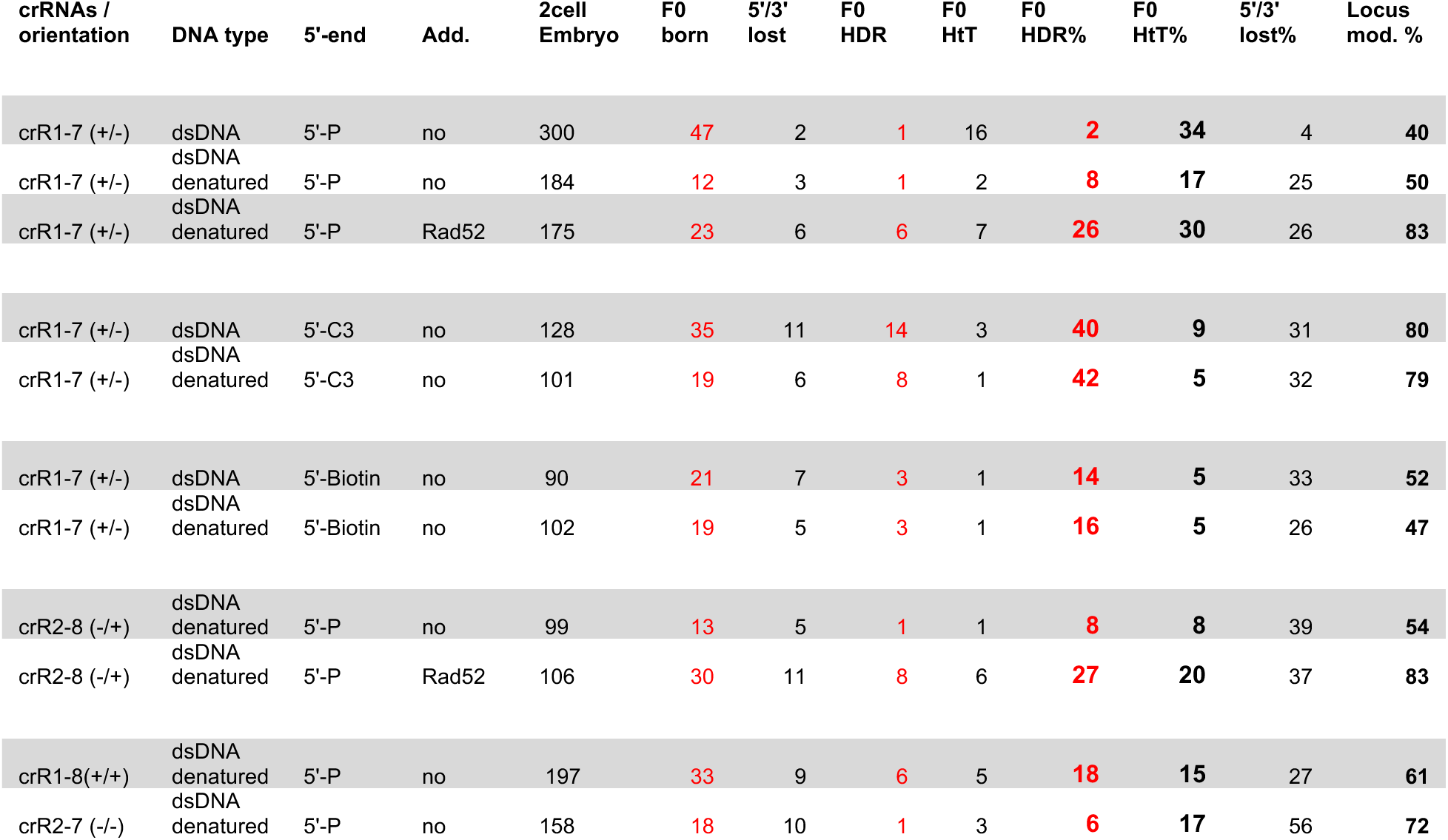
Summary of CRISPR-Cas9 Genome Editing of the *Nup93* Locus. summarizes the results of various experiments targeting the *Nup93* locus using CRISPR-Cas9. The **crRNA/Orientation** column indicates the crRNA pair used and its orientation relative to the *Nup93* gene annotation, where “+” denotes crRNA targeting the antisense (transcriptionally active, template) strand, and “-” denotes crRNA targeting the sense (non-template) strand. **DNA Type** specifies whether double-stranded (dsDNA) or denatured template was used in the respective experiments. The **5’-End** column describes modifications to the 5’ end of the DNA template: “5’-P” denotes a monophosphorylated DNA, “5’-C3” indicates a propyl modification, and “5’-Biotin” refers to a biotin-modified template. **2-cell Embryo** represents the number of 2-cell embryos transferred into recipient mice, while **F0 Born** indicates the number of animals born following the embryo transfer procedure. **5’/3’ Lost** refers to the number of animals with aberrant template insertions at the *Nup93* locus, where either the 5’- or 3’-end of the template was not detected after integration. **F0 HDR** denotes the number of animals with the correctly targeted *Nup93* locus through homology-directed repair (HDR), and **F0 HtT** represents the number of animals showing head-to-tail integration of the donor DNA into the *Nup93* locus. **F0 HDR %** corresponds to the percentage of animals with correct HDR-mediated integration relative to the total number of animals born, while **F0 HtT %** represents the percentage of animals with head-to-tail integration. **5’/3’ Lost %** indicates the percentage of animals with aberrant integrations where the 5’- or 3’-end of the template was not integrated. Finally, **Locus Mod. %** represents the overall percentage of locus modifications observed in each targeting experiment.

The donor DNA template, approximately 600 bp in length (Supplementary Data 2), was designed to include exon-intronic regions flanked by *LoxP* sites (replacing crRNA-targeted regions) and short homologous arms of 60 and 58 nucleotides, respectively. To facilitate the detection of single-copy integrations via Southern blot analysis, we incorporated *Eco*RI and *Bam*HI restriction enzyme recognition sites adjacent to the introduced *LoxP* sequences (Figure 1B, C). To enable efficient targeting, we designed two overlapping crRNAs for each flanking region, targeting complementary DNA strands and exhibiting comparable predicted efficiencies *in silico* (Figure 2, Supplementary Data 1).

### Enhancing HDR with denatured DNA template

Initially, we used crRNA1 and crRNA7, targeting the antisense and sense strands of exon 9 flanking introns, respectively, in combination with double-stranded DNA (dsDNA) templates (Figure 2, 3A). Among 47 pups born, only one exhibited the correctly targeted allele (2%), whereas 16 animals (34%) showed head-to-tail integration of the donor DNA template (Table 1). Consistent with our previous observations, we noted a high frequency of locus modification in 40% of the obtained animals, with 4% carrying insertions of a 5’- or 3’-end degraded donor template (Table 1). To mitigate template multiplication and enhance precise HDR-mediated insertion, we decided to denature dsDNA template for generation of single-stranded DNA (ssDNA) templates for microinjections (Figure 3A). Targeting the *Nup93* locus with denatured donor DNA resulted in a nearly fourfold increase in correctly targeted animals (8%) and an almost twofold reduction in template multiplication (17%) (Table 1). Modification of the locus was observed in 50% of cases, with a notable increase in aberrant template integration, reaching 25% (Table 1). Interestingly, when human RAD52 protein was added to the injection mix containing the denatured DNA template, 26% of the generated animals carried the precise HDR-mediated targeted locus. This represents a more than threefold increase compared to ssDNA alone (and a 13-fold increase compared to dsDNA) in the correct modification of the *Nup93* locus. However, this improvement was accompanied by an unexpected near twofold increase in template multiplication (30%) (Table 1, Figure 4), with approximately 26% of the targeted loci containing a partially degraded template. Overall, locus modification nearly doubled, reaching 83% of the obtained animals (Table 1). These findings underscore the benefits of denatured DNA templates in enhancing HDR efficiency while minimizing template concatemerization in CRISPR/Cas9 genome editing.

**Figure 3.**
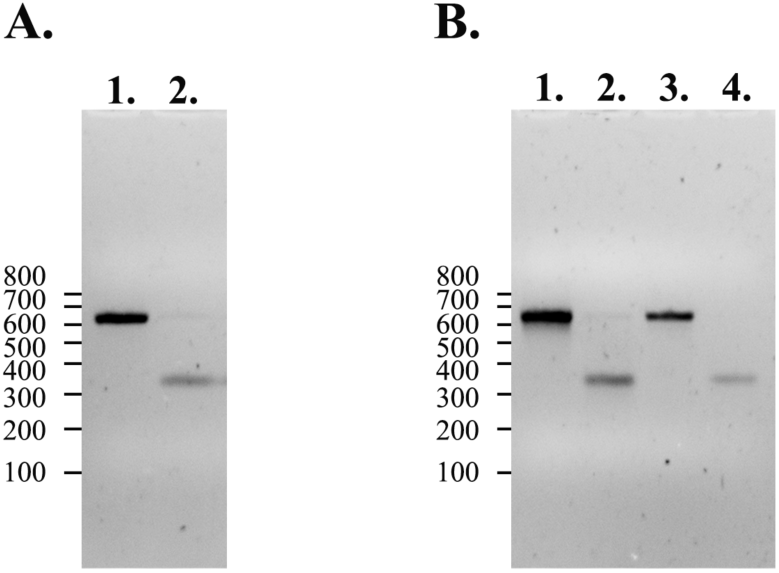
Gel electrophoresis analysis of donor DNA template strandness. Samples were analysed on a 1.5% agarose gel in 1× TAE buffer with ethidium bromide added directly to the gel. DNA was mixed with loading buffer to final concentrations of 0.6 mM HEPES-KOH (pH 7.5), 2 mM potassium acetate, 5% (v/v) glycerol, 0.04% (w/v) bromophenol blue, and 0.04% (w/v) xylene cyanol FF. (**A**) Donor templates digested from plasmid: lane 1, dsDNA; lane 2, denatured DNA. (**B**) 5′-end–modified templates: lane 1, dsDNA with 5′-C3-propyl; lane 2, denatured with 5′-C3-propyl; lane 3, dsDNA with 5′-biotin; lane 4, denatured with 5′-biotin. Size markers (lines) correspond to the GeneRuler DNA Ladder Mix (SM0331, Thermo Fisher Scientific).

**Figure 4.**
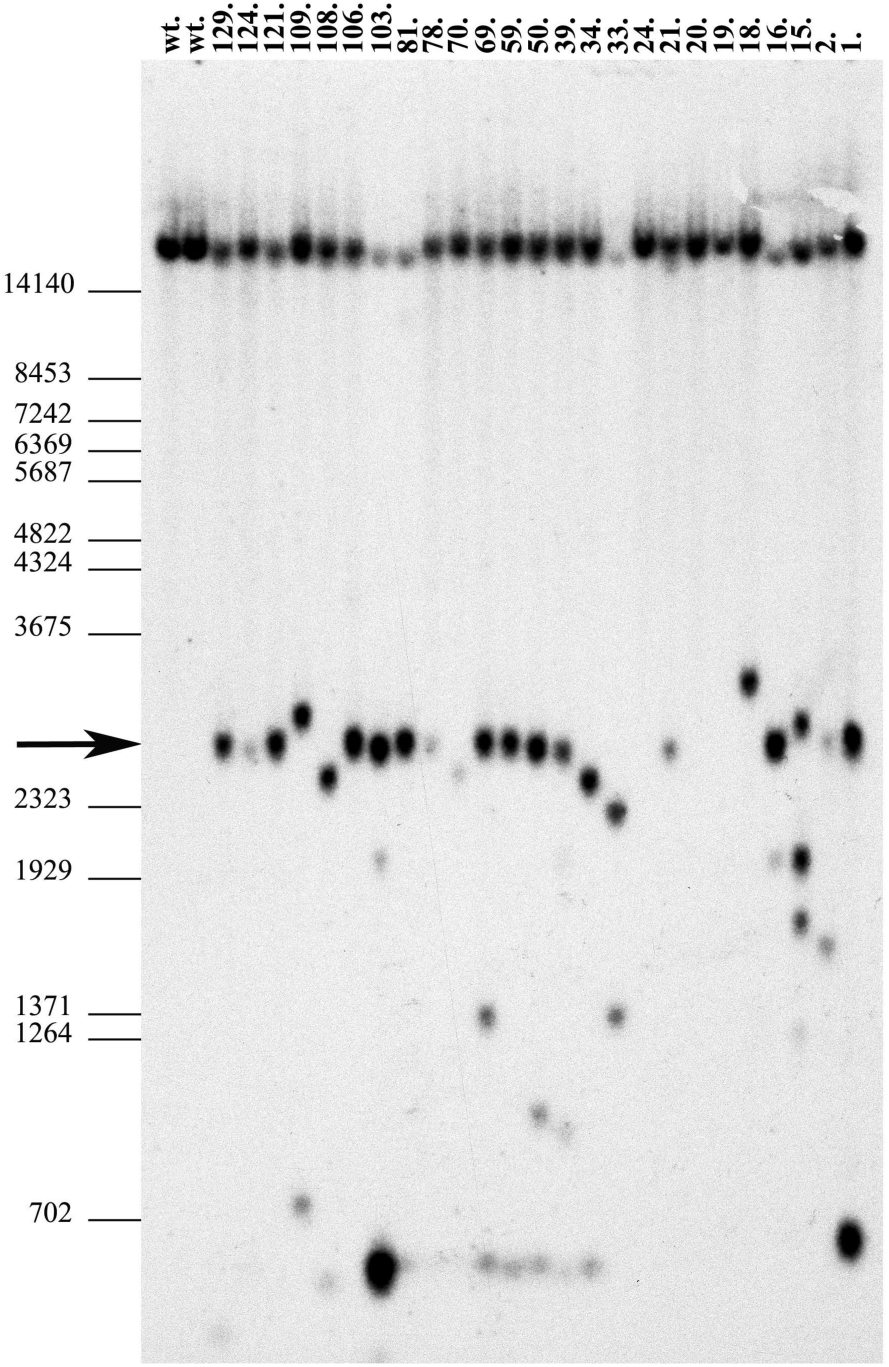
Southern blot analysis of Nup93 locus targeting. DNA samples were extracted from mouse-tail biopsies of F0 offspring and digested with *BamHI*. The following groups of samples were analyzed: Samples 1, 2, 112–129: Microinjected with denatured DNA template and crRNA1–crRNA7 pair. Samples 15–34: Microinjected with denatured DNA template, crRNA1–crRNA7, and Rad52. Sample 39: Microinjected with denatured DNA template and crRNA2– crRNA8 pair. Samples 50–78: Microinjected with denatured DNA template, crRNA2–crRNA8, and Rad52. Samples 79–111: Microinjected with denatured DNA template and crRNA1–crRNA8. Southern blot analysis was performed using a specific probe (Figure 1) following *BamHI* digestion. This detected the wild-type (*wt*) allele (21.7 kb) and two fragments (2.8 kb and 19.0 kb) corresponding to the correctly targeted allele. Notably, under the given conditions, the 21.7 kb and 19.0 kb fragments were not resolved on agarose gels (see *wt* control animals). The presence of a single 2.8 kb band in DNA samples 21, 106, and 121 indicates successful targeting of the *Nup93* allele (*Nup93+/-*). Size marker positions (in bp) are shown on the left. All other animals, except 19, 20, and 24, which contained an unmodified *Nup93* locus, exhibited either head-to-tail or aberrant integrations of the DNA template.

### Strand-specific effects of crRNAs on HDR-mediated integration

Next, we assessed the efficiency of denatured donor DNA template integration using different combinations of selected crRNAs (Table 1). When the crRNA2 and crRNA8 pair, targeting the sense and antisense strands of exon 9 flanking introns, respectively, was used, we observed a similar rate of precise HDR-mediated insertion as with crRNA1 and crRNA7, with approximately 8% of animals carrying the correctly targeted *Nup93* locus (Table 1). Notably, despite a comparable overall locus modification rate (54%), head-to-tail template multiplication was reduced by half (8%) (Table 1). Once RAD52 protein was added to the injection mix, we observed a more than threefold increase in precise HDR-mediated insertion, consistent with the previous crRNA pair. This was accompanied by a threefold increase in animals exhibiting head-to-tail donor DNA multiplication at the modified *Nup93* locus, along with an overall increase in locus modification to 83% (Table 1).

Further analysis of crRNA strand targeting revealed intriguing differences (Figure 2, Table 1). When the crRNA1 and crRNA8 pair, targeting the antisense strand of the *Nup93* region, was used, we detected an almost 2.5-fold increase in precise HDR-mediated insertion of the denatured donor DNA template compared to all other crRNA combinations. Despite this improvement, template multiplication (15%), degradation (27%), and overall locus modification rates (60%) remained similar to other combinations (Table 1, Figure 4). In contrast, targeting the sense strand with the crRNA2 and crRNA7 pair led to an increase in overall locus modification frequency to 72%. However, this increase was driven by aberrant template integration (56%), with only 6% of animals exhibiting correct locus targeting (Table 1). Our findings suggest that crRNA strand selection may influence HDR efficiency and potentially affect the integrity of the denatured donor template during CRISPR/Cas9-mediated targeting of the *Nup93* locus. Targeting the antisense strand, particularly with the crRNA1 and crRNA8 combination, enhances precise HDR-mediated insertion, whereas sense-strand targeting increases overall modification rates but leads to a higher frequency of aberrant integrations (Table 1).

### Impact of 5’-end modifications on genome editing precision

To further optimize our genome editing protocol and assess the impact of 5’-end modified donor DNA templates on reducing head-to-tail concatemers while enhancing HDR-mediated integration, we selected crRNA1 and crRNA7. This pair was chosen based on prior data confirming successful modification of the Nup93 locus using both dsDNA and denatured DNA templates.

In our previous research, we utilized 5’-end C3-spacer (or C3-propyl spacer) modified adapters to prevent ligation events that lead to concatemerization of cDNA molecules [28]. Recent studies in Medaka zygotes have shown that 5’-end C3-spacer and 5’-end C3-biotin modifications can reduce template multimerization [23, 24]. Moreover, recent study in mice showed efficient integration of 5’-biotinilated long DNA templates [25]. Based on these findings, we conducted experiments to evaluate the effects of these modifications on the efficiency of HDR-mediated integration at the *Nup93* locus.

Using a 5’-end biotinylated dsDNA donor template (Figure 3B), we observed a nearly sevenfold increase in HDR-mediated precise locus modification, reaching 14%, compared to just 2% with an unmodified (5’-monophosphorylated) dsDNA template. Additionally, head-to-tail insertions were reduced nearly sevenfold, occurring in only 5% of modified animals (Table 1, Supplementary Figure 2). Despite this improvement, the overall frequency of locus modification remained comparable at approximately 52%. However, an unexpectedly high proportion (33%) of donor templates exhibiting 5’-end or 3’-end degradation upon integration was observed (Table 1). Similar results were obtained when the 5’-end biotinylated dsDNA was denatured prior to microinjection experiments (Table 1).

Application of a donor dsDNA template containing a 5’-end C3-spacer (Figure 3B) resulted in a substantial, nearly 20-fold increase in HDR-mediated insertions, with 40% of modified animals carrying the correct locus modification (Table 1. Supplementary Figure 1). The overall locus modification rate reached approximately 80%, comparable to the rate observed when denatured DNA templates were used in combination with RAD52 protein, but significantly higher than that achieved with the 5’-end biotinylated template (Table 1). However, aberrant insertions involving 5’-end or 3’-end degradation remained frequent, occurring in approximately 31% of cases. Head-to-tail multiple insertions were reduced nearly fourfold, occurring in 8% of animals compared to the unmodified dsDNA template. Denaturation of the 5’-end C3-spacer modified donor template (Figure 3B) did not result in a significant difference compared to the undenatured template, though a slight reduction in head-to-tail concatemerized insertions was observed (5%) (Table 1). Our findings demonstrate that 5’-end modifications, particularly C3-spacer modification, significantly enhance HDR-mediated integration while reducing head-to-tail concatemers.

## Discussion

Historically, genome editing was performed in mouse embryonic stem (ES) cells using specially designed targeting plasmid DNA vectors with large (up to 8kb) homologous arms and positive selection gene [3]. These constructs were carefully verified by sequence analysis before genome integration. Positively selected ES cells were then injected into blastocysts to generate chimeric mice [29]. Achieving germline transmission required extensive breeding of high-percentage chimeras, significantly increasing the number of mice used in experiments. This approach, although effective, was time-consuming and required extensive selection and screening processes. The development of direct CRISPR/Cas9-mediated genome editing in mouse zygotes significantly accelerated the generation of genetically modified mouse models. However, this advancement also introduced new challenges, particularly in the selection of an optimal donor DNA template for precise knock-in modifications. Typically, linear dsDNA donor templates excised from plasmid vectors are preferred due to their ease of generation and the ability to verify sequence integrity before injection. While dsDNA donor templates facilitate precise genome editing in CRISPR/Cas9-mediated knock-in experiments, their use associated with several limitations. One of the major challenges is their low integration efficiency via the homology-directed repair pathway, which is often overshadowed by the more efficient but error-prone non-homologous end joining pathway. Furthermore, dsDNA templates frequently undergo concatemerization and integrate as head-to-tail tandem repeats, complicating precise genome modifications and potentially disrupting endogenous gene regulation [22, 30]. Single-stranded DNA templates have become a powerful tool for increasing the efficiency and precision of CRISPR-Cas9 genome editing [19, 20]. Compared to dsDNA, ssDNA—particularly single-stranded oligodeoxynucleotides (ssODNs)—exhibits higher rates of HDR, making it ideal for introducing precise edits such as point mutations, small insertions, or deletions [19]. However, long ssDNA templates, whether produced enzymatically (e.g., reverse transcription, asymmetric PCR) or through chemical synthesis, often contain errors such as deletions, insertions, or mismatches, depending on the synthesis method and template sequence and length [31–34]. While these templates are generally more efficient than dsDNA, their inherent error rate poses significant risks in applications such as generating mouse models. In some cases, only mice carrying incorrect insertions, point mutations, or unintended deletions caused by synthesis errors are identified among the targeted animals [34] (Skryabin et al., unpublished). To overcome these limitations, we developed a simple and effective method for generating single-stranded DNA (ssDNA) templates by heat denaturation and rapid freezing in liquid nitrogen of dsDNA before adding it to the microinjection mix. Additionally, the use of a low-salt injection buffer (0.6 mM HEPES-KOH, pH 7.5, and 2 mM potassium acetate) allowed to maintain DNA strands in their single-stranded form (Figure 3) while still permitting electropulse-induced microinjection, ensuring efficient delivery into zygotes [35]. This recently developed microinjection method consistently achieved high embryo survival rates, with a geometric mean of 87% across all experiments reported here (Table 2). This indicates that the concentrations of components used in CRISPR-Cas9 editing of the *Nup93* locus were well tolerated by embryos, ensuring that experimental outcomes were not compromised by embryo viability (Table 2) [35].

**Table 2.** Embryo Survival and Transfer Efficiency for *Nup93* Locus Targeting. This table summarizes embryo survival and transfer efficiency for each CRISPR-Cas9 targeting attempt at the *Nup93* locus. The **DNA Template** column specifies whether a double-stranded (dsDNA) or denatured DNA template was used. The **crRNAs/Orientation** column indicates the crRNA pair used and its orientation relative to the *Nup93* gene annotation. The **5’-End -** column describes modifications at the 5’ end of the DNA template, where 5’-P denotes monophosphorylated DNA, 5’-C3 indicates a propyl modification, and 5’-Biotin refers to a biotin-modified template. **Zygotes injected** represents the number of zygotes microinjected with CRISPR-Cas9 components, while **2-Cell survived** indicates the number of zygotes that successfully developed into the 2-cell stage after microinjection. **% Survival** represents the percentage of injected zygotes that developed to the 2-cell stage, reflecting embryo viability after microinjection. **2-Cell Transferred** denotes the number of viable 2-cell embryos transferred into recipient mice. **F0 Born** refers to the number of animals born following the embryo transfer procedure. Finally, **Embryo transf. (%)** represents embryo transfer efficiency as overall percentage of successfully transferred embryos that resulted in live births.

| DNA template | crRNAs (orientation) | 5'-end | Zygotes injected | 2 cells survived | % survivals | 2 cell transferred | F0 born | Embryo transf. % |
| --- | --- | --- | --- | --- | --- | --- | --- | --- |
| dsDNA | crR1-7 (+/-) | 5'-P | 344 | 323 | 94 | 300 | 47 | 16 |
| Denatured dsDNA | crR1-7 (+/-) | 5'-P | 241 | 184 | 76 | 184 | 12 | 7 |
| Denatured dsDNA /Rad52 | crR1-7 (+/-) | 5'-P | 214 | 175 | 82 | 175 | 23 | 13 |
| dsDNA | crR1-7 (+/-) | 5'-C3 | 141 | 128 | 91 | 128 | 35 | 27 |
| Denatured dsDNA | crR1-7 (+/-) | 5'-C3 | 125 | 101 | 81 | 101 | 19 | 19 |
| dsDNA | crR1-7 (+/-) | 5'-Biotin | 93 | 90 | 97 | 90 | 21 | 23 |
| Denatured dsDNA | crR1-7 (+/-) | 5'-Biotin | 108 | 102 | 94 | 102 | 19 | 19 |
| Denatured dsDNA | crR2-8 (-/+) | 5'-P | 199 | 179 | 90 | 99 | 13 | 13 |
| Denatured dsDNA /Rad52 | crR2-8 (-/+) | 5'-P | 153 | 136 | 89 | 106 | 30 | 28 |
| Denatured dsDNA | crR1-8 (+/+) | 5'-P | 228 | 197 | 86 | 197 | 33 | 17 |
| Denatured dsDNA | crR2-7 (-/-) | 5'-P | 212 | 158 | 75 | 158 | 18 | 11 |
|  |  |  | Total: 2058 | Total: 1773 | Mean: 87% | Total: 1640 | Total: 270 | Mean: 16,3% |

Consistent with previous reports on ssDNA application, we observed a four- to ninefold increase in overall HDR-mediated efficiency for precise *Nup93* locus targeting, along with an almost twofold reduction in head-to-tail template integration (Table 1, Figure 5). The key advantage of our proposed method is its flexibility in targeting construct design, as it is not constrained by structural or sequence-related complications. Furthermore, the linear DNA template, digested from plasmid vectors, maintains sequence integrity, and the denaturation step enables the introduction of both ssDNA strands for HDR. As a result, any ssDNA strand-dependent effects can be disregarded [36, 37]. Interestingly, previous cell culture studies have suggested that CRISPR- Cas9-mediated genome editing rates (mutation frequencies via the NHEJ mechanism) depend on the targeted DNA strand. When crRNAs (guide RNAs) were designed to target transcriptionally active template strands (antisense strands), they were associated with higher mutagenesis rates compared to crRNAs complementary to the non-template (sense) strands [38]. However, our *in vivo* results revealed almost no difference in *Nup93* locus modification frequencies between crRNA pairs targeting the template and non-template strands, with rates of 61% and 72%, respectively (Figure 2, Table 1). Notably, Nup93 is an essential component of the nuclear pore complex, expressed in mouse zygotes (SRP086707), and its depletion leads to severe developmental defects, underscoring its crucial role in early embryogenesis [39–41]. Interestingly, our results demonstrate that transcriptionally active - template-strand (antisense-strand) targeting led to a significant, nearly 2.5-fold increase in HDR-mediated precise editing of the Nup93 locus, compared to non-template and other crRNA combinations (Table 1, Figure 2, 5). These findings offer valuable insight into strand-dependent effects on CRISPR-Cas9 mediated targeting and HDR efficiency *in vivo* and suggest a potential tempting strategy for optimizing precise genome editing, particularly for genes that are transcriptionally active during early embryonic development.

**Figure 5.**
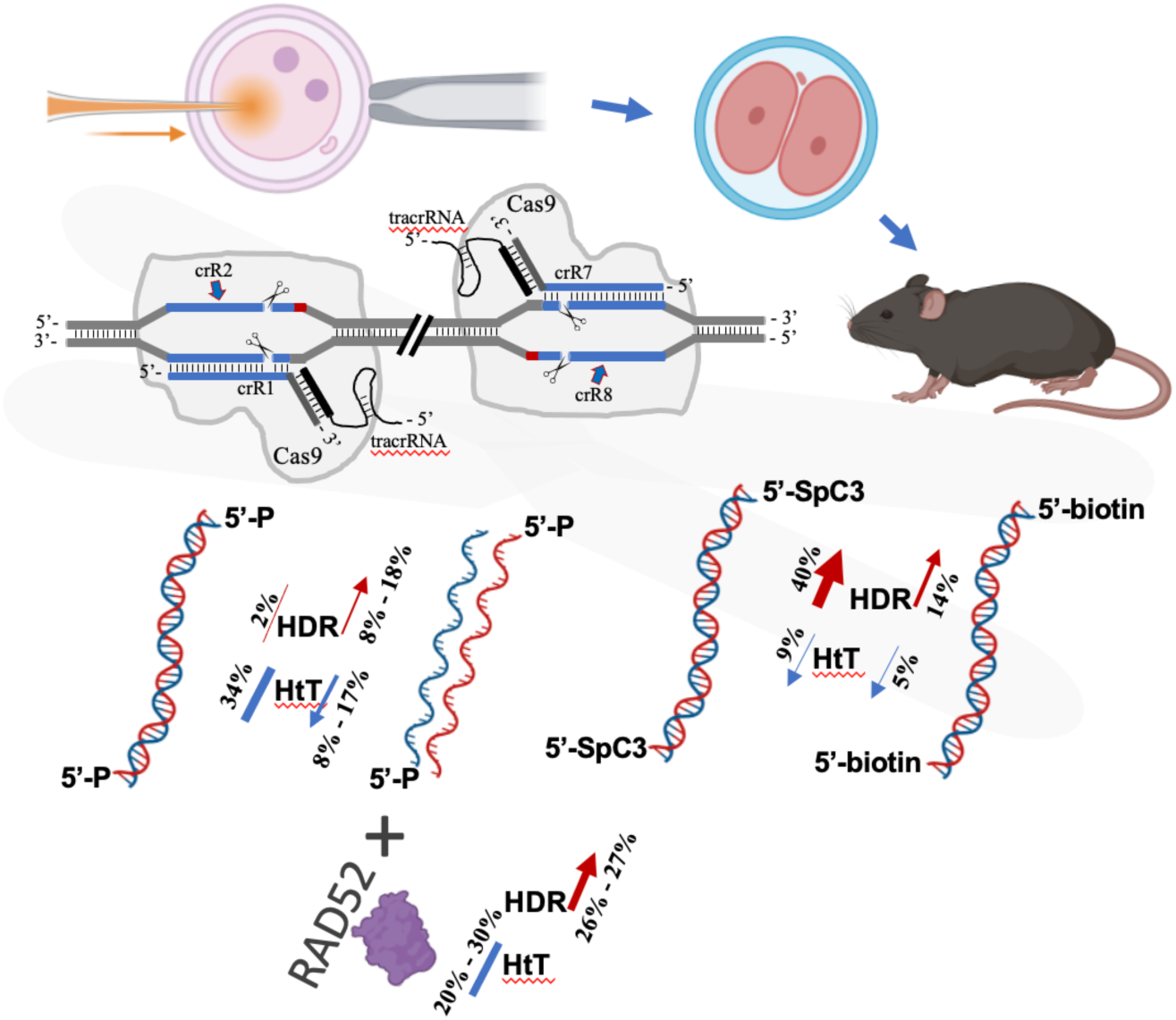
Schematic summary of Nup93 locus targeting.

Another interesting finding came from an application of human recombinant RAD52 protein in CRISPR-Cas9 mediated knock-in targeting of *Nup93* locus with denatured donor DNA template. Human RAD52 is an essential protein for DNA repair process, it facilitates the annealing of complementary single-stranded DNA and thereby maintaining genomic stability [18, 42]. Previous studies in mammalian system implicating RAD52 into enhancing HDR efficiency were performed in cell culture (*ex vivo*) using expression vectors encoding human protein, Cas9-RAD52 fusion or Rad52 yeast homolog [43–45]. To the best of our knowledge, this study represents the first direct application of human RAD52 protein during genome editing of mouse zygotes. Strikingly, we observed a notable increase in overall locus targeting efficiency, with over 80% of obtained animals harbouring modifications in the *Nup93* locus (Table 1). Furthermore, HDR efficiency showed a more than threefold enhancement in precise knock-in events. On the other hand, we also detected an approximately twofold increase in head-to-tail tandem integration of the donor DNA template and similar around 30% rate of degraded template integration (Table 1), suggesting that a number of multiplication events occur using HDR mechanism [30]. These findings underscore the potential of RAD52 as a potent enhancer of HDR-based genome editing strategies in mouse zygotes. Notably, the use of DNA templates with 5′-end modifications that block the phosphate group could potentially further optimize RAD52-mediated enhancement, preventing template concatemerization improving the precision and efficiency of targeted genome modifications.

Here, we also investigated HDR efficiency using donor DNA template containing 5’-biotin and 5’-propyl spacer modifications. Previous studies have shown that modifying the donor DNA ends can significantly enhance knock-in efficiency in mammalian cells [23, 26]. Additionally, a study showed that 5′ biotinylation, in combination with a Cas9-Streptavidin fusion protein (Cas9-mSA), improved targeted integration by up to fourfold in mouse embryos by guiding the donor DNA to the DSB site [27]. More recent findings in mouse embryos suggest that 5’ biotin modifications alone can enhance single-copy HDR, even in the absence of a Streptavidin-coupled Cas9 [25]. In this study, we applied a 5′-end biotinylated DNA donor templates and observed a nearly sevenfold increase in HDR-mediated precise locus modification compared to an unmodified dsDNA template, reaching 14% to 16% efficiencies (Table 1). However, when compared to a denatured unmodified DNA template used with the same crRNA pair, the increase in HDR efficiency was only about twofold (Table 1). Notably, the beneficial effects of 5′-end biotinylation were diminished when compared to results obtained from using a combination of crRNA1 and crRNA8 (targeting the template strand) along with a denatured DNA template (Table 1). One notable advantage of 5′-biotinylation was its ability to significantly reduce head-to-tail template multiplication. However, this advantage must be weighed against potential drawbacks, including the risk of PCR-amplified or chemical synthesis errors, which can introduce unintended genetic variations. This trade-off highlights the importance of selecting the most suitable donor template modification strategy based on the specific requirements of a given experiment. In the context of *Nup93* locus modification, the benefits of 5′-end biotinylation were not as pronounced when compared to the use of unmodified denatured DNA templates.

In addition to 5′-biotinylation, we explored the effects of a 5′-end C3-spacer modification on HDR efficiency. Remarkably, the use of a 5′-C3 spacer-modified dsDNA donor template resulted in an almost 20-fold increase in HDR-mediated insertions compared to an unmodified dsDNA template and a nearly fivefold increase compared to a denatured unmodified template. This finding suggests that the C3 spacer modification substantially enhances HDR efficiency, likely by altering the stability and/or accessibility of the donor template at the repair site. Similar to 5′-biotin modification, the single-copy HDR integration efficiency was comparable between 5′-C3 spacer-modified, both denatured DNA and dsDNA templates, indicating that this modification facilitates precise and efficient repair in both single-stranded and double-stranded donor contexts (Table 1). Interestingly, an increase in overall locus targeting efficiency was observed, with approximately 80% of obtained animals carrying modifications at the *Nup93* locus. This efficiency was similar to that achieved with RAD52-mediated HDR enhancement and unmodified denatured DNA template utilization. However, a key distinction was the drastic reduction in head-to-tail multiplication of donor templates, a feature comparable to the effect observed with 5′-biotinylated templates. This suggests that the 5′-C3 spacer modification may serve as an effective alternative strategy for reducing concatemer formation while maintaining high HDR efficiency, making the C3 spacer modification a compelling strategy for precise genome editing.

Although this study was conducted at a single genomic locus, it provided a well-controlled and systematic approach to evaluating factors that enhance HDR-mediated precise genome editing (Figure 5). By carefully modifying only one parameter at a time—such as denaturation and 5′-end modifications of the donor DNA template, or the presence of RAD52—we were able to precisely assess the direct impact of each modification on HDR efficiency and overall locus targeting. This meticulous approach minimized potential confounding variables, as all comparisons were performed under identical experimental conditions using the same crRNA pair. In cases where strand targeting were analysed, we ensured consistency by using identical donor templates with different crRNA pairs.

The controlled nature of our study provides a solid basis for understanding the potential benefits of these strategies and their role in optimizing genome editing outcomes. While our findings demonstrate significant improvements in HDR efficiency, additional studies across multiple genomic loci are necessary to validate the broader applicability of these approaches. Nevertheless, we anticipate that while locus-specific variations may occur, the overall trends observed in this study are likely to be reproducible across different genomic loci. These insights offer valuable guidance for refining CRISPR-Cas9-mediated genome editing strategies, ultimately contributing to more efficient and precise genetic modifications in mammalian models

## Conclusions

Our findings highlight key advancements in optimizing CRISPR-Cas9-mediated HDR efficiency in mouse zygotes. The use of heat-denatured DNA templates significantly enhances precise genome modifications while reducing unwanted concatemerization. A major advantage of denatured templates is their complete sequence fidelity, as they are derived directly from verified double-stranded DNA, ensuring the absence of synthesis-induced errors. Additionally, RAD52 supplementation serves as a potent enhancer of HDR efficiency, although its use should be carefully balanced against increased template multiplication. Moreover, 5′-biotinylation and 5′-C3 spacer modifications present promising strategies for improving HDR-based genome editing. While biotinylation effectively reduces template concatenation, C3 spacer modifications provide a striking improvement in HDR efficiency while maintaining precise genome integration. These findings suggest that modifying donor templates while targeting specific gene loci can substantially improve CRISPR-based genome editing outcomes. However, a potential drawback of using 5′-modified templates is the risk of synthesis errors, which may introduce unintended sequence variations. We believe that these considerations will guide researchers in selecting the most robust approach for generating mouse models. The choice ultimately depends on balancing high HDR efficiency with a low but existing risk of errors versus prioritizing absolute sequence integrity at the potential cost of reduced efficiency. Future studies should investigate the combinatorial effects of these modifications and their interactions with other HDR-enhancing factors to further refine genome editing strategies. Additionally, further investigation into transcriptionally active strand targeting may provide valuable insights for optimizing precise genome editing strategies. Collectively, these advancements represent important steps towards improving the efficiency, precision, and reliability of CRISPR/Cas9-mediated genome engineering in mammalian systems.

## MATERIALS AND METHODS

### Generation of the *Nup93*-flox mice

*Nup93*-flox mouse line was generated using “one-step” strategy for the generation of cKO mouse models by direct oocyte microinjections of CRISPR/Cas9 complexes and DNA template, containing two *LoxP* sites (*Nup93*-template) (Supplementary Data 2). Donor DNA template (*Nup93*-template) for microinjection was chemically synthesized and cloned into pUC57 vector [Biomatik, USA]. DNA template was cut out directly from plasmid, or PCR amplified from the aforementioned plasmid vector with 5’-end C3-spacer (C3-propyl) or 5’-end Biotin modified direct NUP93dB and reverse NUP93rB primers. The resulting dsDNA fragments were purified using 1% agarose gel electrophoresis, extracted with 6 M NaI, and subsequently stored in ddH2O. For the preparation of CRISPR/Cas9 microinjection solution, commercially synthesized crRNA’s: Nup_crRNA1 or Nup_crRNA2 targeting the left flank, and Nup_crRNA7 or Nup_crRNA8 targeting the right flank, were mixed with tracrRNA (Integrated DNA Technologies (IDT), USA) in 10 mM potassium acetate, 3 mM Hepes-KOH (pH 7.5) buffer (Table 1, Figure 2, Supplementary Data 1). The mixture was incubated at 95 °C for 2 min, followed by cooling to room temperature to allow annealing. The resulted crRNA/tracrRNA complexes were combined with *in vitro* transcribed Cas9 mRNA and Cas9 protein (Alt-R *S.p*. Cas9, IDT, USA). The dsDNA *Nup93* templates were either used directly or were denatured at 95 °C for 5 minutes, rapidly cooled in liquid nitrogen, and then added to the microinjection mix. The final microinjection buffer (0.6 mM Hepes-KOH (pH 7.5) and 2 mM potassium acetate) contained the following CRISPR-Cas9 components: crRNAs (1 pmol/µl each), tracrRNA (2 pmol/µl), Cas9 mRNA (20 ng/µl), Cas9 protein (40 ng/µl), *Nup93* template (0.025 pmol/µl). When human recombinant RAD52 protein (Abcam, ab188614) was included, its final concentration in the solution was 0.05 pmol/µl. The injection solution was filtered through Millipore centrifugal columns and spun at 10,000g for 10 min at room temperature. Microinjections were performed in B6D2F1 zygotes, a hybrid of C57BL/6J and DBA mouse strains. Cytoplasmic microinjections were conducted in M2 media using the Transjector 5246 (Eppendorf) and Narishige NT-88NE micromanipulators, mounted on a Nikon Diaphot 300 inverted microscope. The procedure utilized our patented electropulse-induced microinjection method with a single electrode, applying the following parameters: injection pressure of 120–130 hPa, pulse amplitude of 85 V, frequency of 100 kHz, impulse package duration of 600 µs, and drilling at 50 packages per second [35]. The following day, two cells embryos that survived microinjections were transferred to oviducts of pseudopregnant CD1 foster mice and carried to term. Positively targeted F0 and F1 animals were identified by qPCR, Southern blot (Figure 4, Supplementary Figure 1-3) and sequencing analysis of genomic DNA isolated from tail biopsies.

### qPCR analysis of the targeting events and sequencing of the *Nup93* exon 9 genomic locus

TaqMan qPCR analysis was performed in 20µl volume using LightCycler 480 system (Roche). For detection of the 5’ and 3’ correctly targeted regions we designed assays with external primers (PDE_d1 and PDE_r1, respectively), located outside of the targeting homology and internal primers LoxA1rev and LoxA2dir, respectively. TaqMan UPD probe #74 (Roche, cat.no. 04688970001) was used in both assays. Hence, 5’ assay was performed with Nup93_d3, LoxA1rev and TaqMan UPD probe: #74 (Roche); whereby for 3’ assay the following primers were used: Nup93_r3, LoxA2dir and TaqMan UPD probe: #74. The correct size (of 254 bp and 270 bp for 5’ and 3’ assays, respectively) of resulting qPCR amplicons were confirmed using 6% (w/v) polyacrylamide gel (1 X TBE buffer) electrophoresis followed by ethidium bromide staining and sequencing. *Nup93* targeted genomic locus was sequenced using PCR primers pairs for 5’ region (736 bp): Nup93_d4 / LoxA2rev, and for 3’ region (714 bp): LoxA1dir / Nup93_r4.

### Southern blot analysis of genomic DNA

PCR-positive F0 and F1 animals were analyzed by Southern blot. Genomic DNA (5–10 µg) was digested with *Bam*HI (for F0 mice) or *Bam*HI/*Eco*RI (for F1 animals), fractionated on 0.8% agarose gels, and transferred onto GeneScreen nylon membranes (NEN DuPont) (Figure 4, Supplementary Figure 1-3). The membranes were hybridized with a ³²P-labeled HR probe and subsequently washed at 65°C with a final concentration of 0.5× SSPE (1× SSPE: 0.18 M NaCl, 10 mM NaH2PO4, 1 mM EDTA, pH 7.7) and 0.5% sodium dodecyl sulfate (SDS).

### Animals used for generation of *Nup93-*flox mouse line

All mouse procedures were performed in compliance with the guidelines for the welfare of experimental animals issued by the Federal Government of Germany. The mouse line was established by breeding positively identified founder animals with C57BL/6J mice to produce heterozygous mice. Pups were weaned at 21 to 28 days after birth, and females were kept separately from males. The mice were housed in standard IVC cages. General health checks were performed regularly in order to ensure that any findings were not the result of deteriorating physical conditions of the animals.

### Programs and Software

Mouse *Nup93* locus analysis and crRNA design were performed using the UCSC BLAT mouse genome browser (mm39 assembly: https://genome.ucsc.edu/cgi-bin/hgBlat), followed by further sequence analysis with SeqBuilder Pro and SeqMan Pro (for analyses of sequencing reads) from the DNASTAR Lasergene 17 package. Tables were created using Microsoft Excel, while Microsoft PowerPoint, Adobe Photoshop, BioRender.com, and Keynote (Apple Inc.) were used for figures generation. English language refinement was partially assisted by ChatGPT 3,5.

### Institutional Review Board Statement

All mouse studies conducted at the Transgenic animals and genetic engineering models (TRAM) core facility of the University Clinic Muenster were performed in compliance with the guidelines for the welfare of experimental animals issued by the Federal Government of Germany and approved by the State Agency for Nature, Environment and Consumer Protection North Rhine-Westphalia LANUV (Landesamt für Natur, Umwelt und Verbraucherschutz Nordrhein-Westfalen) (protocol code: AZ: 84-02.04.2016.A546).

## Author Contributions

B.V.S., D.A.B., H.P., and T.S.R. conceived the project. B.V.S. and T.S.R designed the study. H.K., L.G., B.S., A.S., B.V.S., and T.S.R. were involved in experimental work. B.V.S. and T.S.R. participated in the preparation of the figures. B.V.S., T.K., and T.S.R. analyzed the data and wrote the paper. All authors have read and agreed to the published version of the manuscript.

## Acknowledgments

The authors would like to thank Nina Usatchewa for her professional and excellent assistance with mouse care.

## Funding

This work was supported by the Deutsche Forschungsgemeinschaft (DFG BR-5733/4-1), and the IZKF Muenster (IZKF-Brau2/013/19). Core Facility TRAM is an institution of the Medical Faculty of the University of Münster. The authors gratefully acknowledge the support of the Medical Faculty.

## Conflicts of Interest

B.V.S., L.G., and T.S.R. are co-inventors of patent WO2022/043355A1. The other co-authors have no conflict of interest to declare.

## Notes

### Competing Interest Statement

The authors have declared no competing interest.

