## Supplementary Data for "CRISPR-Cas9 HDR Optimization: RAD52, Denatured and 5’-Modified DNA Templates in Knock-In Mice Generation"

**Running head: Highly Efficient Mouse Genome Editing**

| | |
|---|---|
| Guide Sequence Nup_crRNA1 | CTGCACTGAATTTGAGCCAG |
| Protospacer Adjacent Motif (PAM) | GGG |
| MIT Guide Specificity Score | 70 |
| Efficiency: Doench et al. 2016 Score (raw) | 58 |
| Efficiency: Doench et al. 2016 Score (percentile) | 75 |
| Efficiency: Moreno-Mateos T7 Score (raw) | 84 |
| Efficiency: Moreno-Mateos T7 Score (percentile) | 98 |
| Efficiency: Doench et al 2014 Score | 16 |
| Bae et al. Out-of-Frame Score | 66 |

| | |
|---|---|
| Guide Sequence Nup_crRNA2 | TCAAATTCAGTGCAGAAGAT |
| Protospacer Adjacent Motif (PAM) | GGG |
| MIT Guide Specificity Score | 65 |
| Efficiency: Doench et al. 2016 Score (raw) | 55 |
| Efficiency: Doench et al. 2016 Score (percentile) | 66 |
| Efficiency: Moreno-Mateos T7 Score (raw) | 36 |
| Efficiency: Moreno-Mateos T7 Score (percentile) | 31 |
| Efficiency: Doench et al 2014 Score | 29 |
| Bae et al. Out-of-Frame Score | 69 |

| | |
|---|---|
| Guide Sequence Nup_crRNA8 | TGTCCGTGTCTGGTCAGGGC |
| Protospacer Adjacent Motif (PAM) | AGG |
| MIT Guide Specificity Score | 74 |
| Efficiency: Doench et al. 2016 Score (raw) | 52 |
| Efficiency: Doench et al. 2016 Score (percentile) | 56 |
| Efficiency: Moreno-Mateos T7 Score (raw) | 43 |
| Efficiency: Moreno-Mateos T7 Score (percentile) | 46 |
| Efficiency: Doench et al 2014 Score | 5 |
| Bae et al. Out-of-Frame Score | 67 |

| | |
|---|---|
| Guide Sequence Nup_crRNA7 | AAGCCTGCCCTGACCAGACA |
| Protospacer Adjacent Motif (PAM) | CGG |
| MIT Guide Specificity Score | 70 |
| Efficiency: Doench et al. 2016 Score (raw) | 60 |
| Efficiency: Doench et al. 2016 Score (percentile) | 81 |
| Efficiency: Moreno-Mateos T7 Score (raw) | 54 |
| Efficiency: Moreno-Mateos T7 Score (percentile) | 69 |
| Efficiency: Doench et al 2014 Score | 29 |
| Bae et al. Out-of-Frame Score | 59 |

**Supplementary Data 1. *In silico* calculated efficiency of crRNAs used in this study.**
The data were obtained from USCS BLAT mouse genome browser (mm39 assembly).

*CAAGCAGGGCAGGGATATACTCATCAGTGATCAGAGCTCTTAGGATTGGCAAGGCAGA*
*GG*AATTCAAGCTGGGCCCTGGTAATATAACTTCGTATAGCATACATTATACGAAGT
TATGGCTGCTGCCCAGCAGGAAACAACCTACTTGAGGGCAACCCTTCCTCATTCTG
CCTGCCAGTGCAGTCATAGCCAGGTAACTCCTCCCCTTGTACATGGTTGTCATACA
CAACCTCATTTATATGCAG**AGCATTTCTGACATGTGGGCCATGGTAAAACAAAT**
**GACAGATGTGGTGCTGACACCAGCAACTGACGCCCTGAAGAGCCGGAGCAGT**
**GTGGAAGTGCGTATGGACTTTGTCAAGCAAGCCTTGGGATATCTTGAGCAGA**
**G**GTAAGGAAGCTCGCTCTCTCGCTCTCTCGCTCTCTCTCTCTCTCTCTCTCTC
TCTCTCTCTCTCTCTCCAGGTTAGGATCGCATAGGATAACTTCGTATAGCATACATT
ATACGAAGTTATGGCTGCTGCCCAGCATCCTTAATGCGCGTAGTCGGATCC*GCAGG*
*CTTTCACCTTCCCTAAAGTCACATGTTACTTGGTCTTACGTAGTTCTGGGACT*

**Supplementary Data 2. DNA template sequence used in the study**

The Nup93 template sequence is shown with homology arms in italics, *Lox*P sites underlined, and exon 9 highlighted in bold and underlined.

**List of PCR primers used in the study:**

| | |
|---|---|
| Nup93_d1 | GCATGGTAATTGGTCTGAAAGCC |
| Nup93_d2 | CCTATAATGATGGAGGTGCTTCTGTC |
| Nup93_d3 | GGGTGAGCCACTAAGGTTGAGC |
| Nup93_d4 | AAGCCCTAAGAAACTTAAATAACCTTGTAA |
| Nup93_r1 | AGAATTAGCTCAGCTCCAGTCAGTTT |
| Nup93_r2 | GTGCAGAAACAGTCTGAGTAGATAAGTG |
| Nup93_r3 | CACCACCCTTTCCATGTATGATTT |
| Nup93_r4 | CGTGTTAAATTATGGAAAGCAGAGAA |
| LoxA1dir | AAGCTGGGCCCTGGTAAT |
| LoxA1rev | GCCCTCAAGTAGGTTGTTTCC |
| LoxA2dir | CCAGGTTAGGATCGCATAGG |
| LoxA2rev | CGACTACGCGCATTAAGGA |

5’-end C3-spacer (C3-propyl) or 5’-end Biotin modified primers:

| | |
|---|---|
| NUP93dB | CAAGCAGGGCAGGGATATACTCAT |
| NUP93rB | AGTCCCAGAACTACGTAAGACCAAGTAAC |

**Supplementary Data 3. List of PCR primers used in this study**

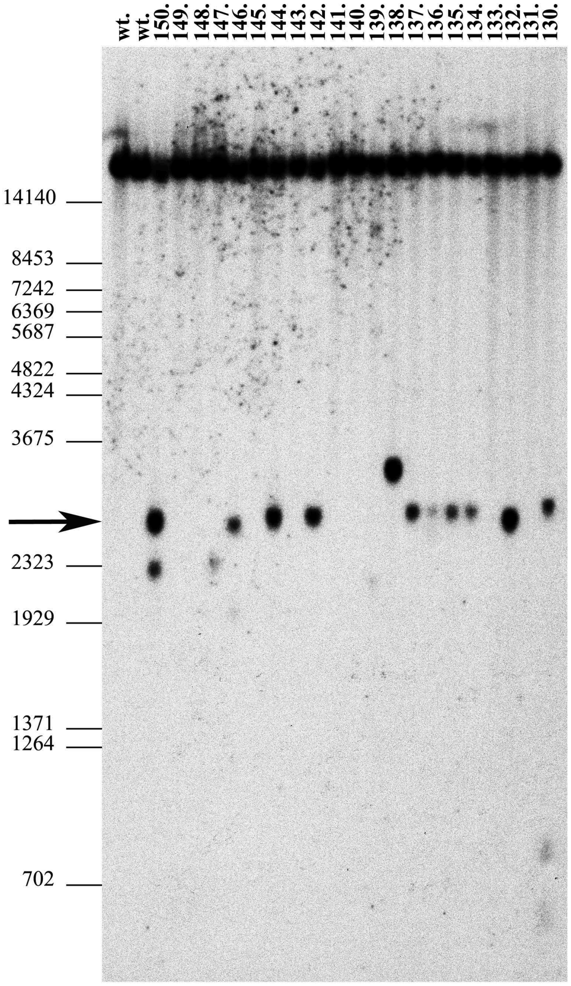


**Supplementary Figure 1. Southern blot analysis of Nup93 locus targeting in FO offsprings**.

DNA samples were extracted from mouse-tail biopsies of F0 offspring and digested with *BamHI*. The following groups of samples were analyzed: Samples 130–150: Microinjected with non-denatured DNA template, labelled 5'C3, and crRNA1–crRNA7 pair. Southern blot analysis was performed using a specific probe (Figure 1) following *BamHI* digestion. This detected the wild-type (*wt*) allele (21.7 kb) and two fragments (2.8 kb and 19.0 kb) corresponding to the correctly targeted allele. Notably, under the given conditions, the 21.7 kb and 19.0 kb fragments were not resolved on agarose gels (see *wt* control animals). The presence of a single 2.8 kb band in DNA samples 134 -137, 142, 144, and 146 indicates successful targeting of the *Nup93* allele. Size marker positions (in bp) are shown on the left.

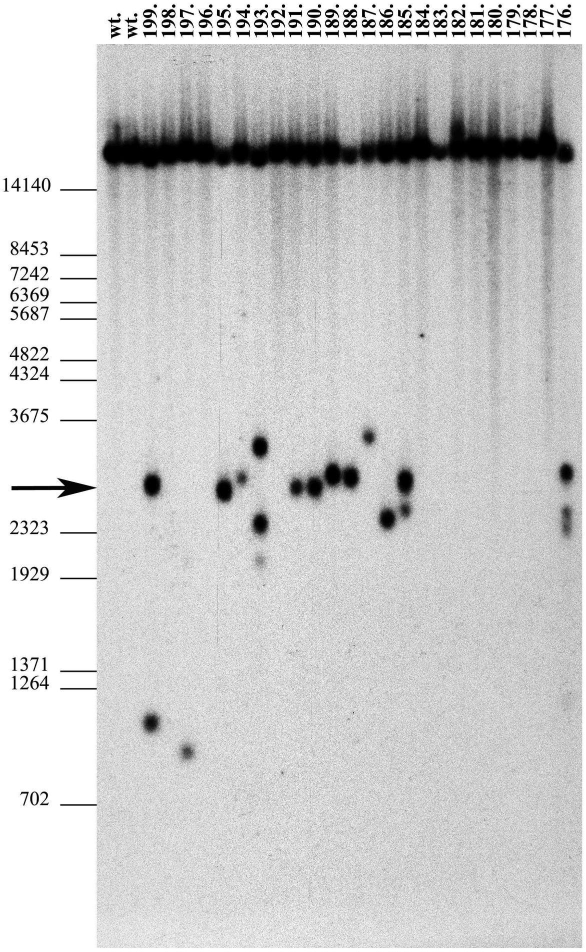


**Supplementary Figure 2. Southern blot analysis of Nup93 locus targeting in FO offsprings**

DNA samples were extracted from mouse-tail biopsies of F0 offspring and digested with *BamHI*. The following groups of samples were analyzed: Samples 176–183: Microinjected with denatured DNA template, labelled 5’C3, and crRNA1–crRNA7 pair. Samples 184–199: Microinjected with non-denatured DNA template, labelled 5’Biotin, and crRNA1–crRNA7 pair. Southern blot analysis was performed the same way, as described in supp. fig. 1. The presence of a single 2.8 kb band in DNA samples 188, 189, and 194 indicates successful targeting of the *Nup93* allele. Size marker positions (in bp) are shown on the left.

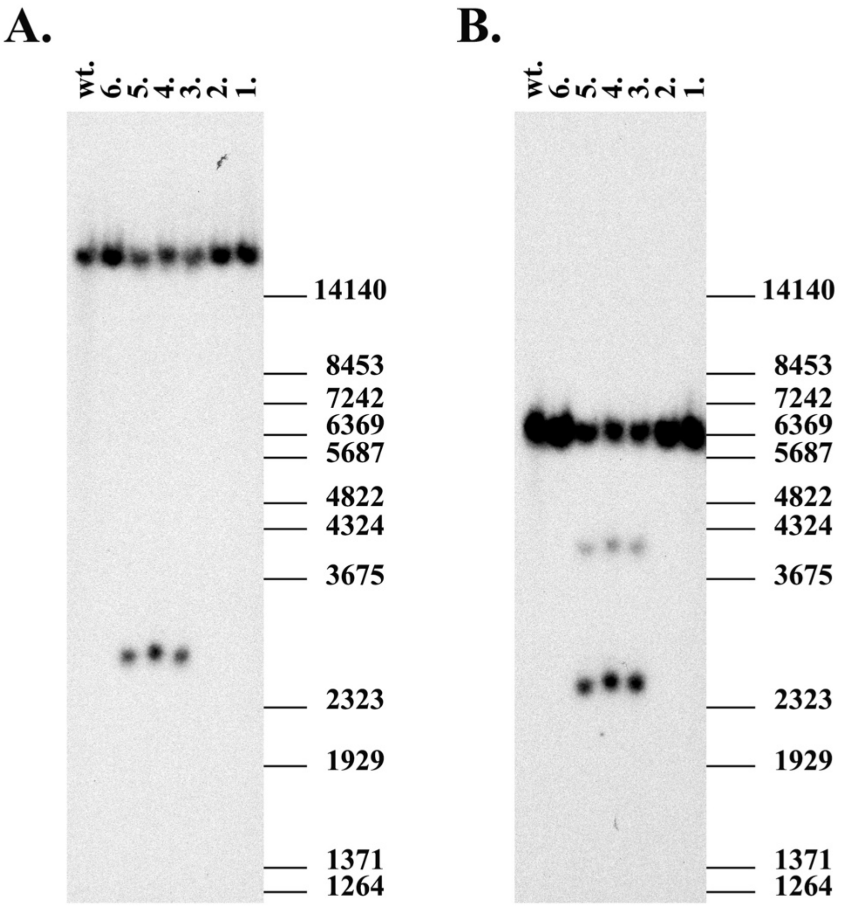


**Supplementary Figure 3**. **Southern blot analysis of F1 offspring**.

Southern blot analysis of genomic DNA isolated from mouse tail biopsy (numbers 1-6, and wt) and hybridized with the Nup93 HR probe. **A**: *Bam*HI enzymatic digestion revealed the wild type allele (21.7 kb) and targeted allele at (2.8 kb). DNA samples obtained from mice numbers 3 - 5 contain correctly targeted *Nup93* allele. Positions of the size marker (in bp) are shown on the right. **B**: *Eco*RI enzymatic digestion, to prove F1 mice with correctly targeted allele. The wild type allele corresponds to band 6.5 kb and targeted allele corresponds to 2.5 kb and 4.0 kb bands. DNA samples 3 - 5 contain correctly targeted *Nup93* allele.
